# Actin activates *Pseudomonas aeruginosa* ExoY nucleotidyl cyclase toxin and ExoY-like effector domains from MARTX toxins

**DOI:** 10.1101/032201

**Authors:** Dorothée Raoux-Barbot, Cosmin Saveanu, Abdelkader Namane, Vasily Ogryzko, Lina Worpenberg, Souad Fellous, Elodie Assayag, Daniel Ladant, Louis Renault, Undine Mechold

## Abstract

*Pseudomonas aeruginosa* is a major cause of chronic infections in cystic fibrosis patients. The nucleotidyl cyclase toxin ExoY is a virulence factors injected by the pathogen and associated with severe damage to lung tissue. ExoY-like cyclases are also found in other Gram-negative pathogens and shown to contribute to virulence, although they remained poorly characterized. Here we demonstrate that filamentous actin (F-actin) is the hitherto unknown co-factor that activates *P. aeruginosa* ExoY within host target cells. Highly purified actin, when polymerized into filaments, potently stimulates (>10,000 fold) ExoY activity. ExoY co-localizes *in vivo* with actin filaments in transfected cells and, *in vitro*, it interferes with the regulation of actin assembly/disassembly-dynamics mediated by important F-actin-binding proteins. We further show that actin also activates an ExoY-like adenylate cyclase from a *Vibrio* species. Our results thus highlight a new sub-class within the class II adenylyl cyclase family, defined as actin-activated nucleotidyl cyclase (AA-NC) toxins.

## INTRODUCTION

*Pseudomonas aeruginosa* is an opportunistic human pathogen that causes severe infections in immune-compromised individuals and is a major cause of chronic infections in cystic fibrosis patients. Equipped with a type III secretion system (T3SS), *P. aeruginosa* can inject effector proteins directly into the host cells where they contribute to virulence of the pathogen [for review see ^1,2^]. Four different T3SS delivered effectors have been characterized (exoenzyme T, Y, U and S), but new effectors were recently identified^3^. Exoenzyme Y (ExoY) is present in 89% of clinical isolates^4^, and was originally identified as an adenylate cyclase in 1998 (ref. ^5^) due to amino acid sequence homology with two well characterized class II adenylate cyclase toxins, CyaA from *Bordetella pertussis* and edema factor (EF) from *Bacillus anthracis*. Recent results revealed that substrate specificity of these enzymes expressed in cell cultures is not restricted to ATP: EF and CyaA were shown to use UTP and CTP as substrate^6^ while ExoY was shown to promote the intracellular accumulation of several cyclic nucleotides^7,8^ with a preference for cGMP and cUMP over cAMP and cCMP formation^7^. The physiological effects of ExoY resulting from accumulation of these cyclic nucleotides include the hyperphosphorylation of tau and the disruption of microtubules causing the formation of gaps between endothelial cells and increased permeability of the endothelial barrier^8,9,10,11^. Most recent results showed that ExoY presence correlates with long term effects on recovery after lung injury from pneumonia^12^.

Recent whole genome sequencing projects have identified ExoY nucleotidyl cyclase modules among a variety of toxic Multifunctional Autoprocessing repeats-in-toxins (MARTX) effector domains in multiple bacterial species of the *Vibrio genus^13^* that represent emerging human or animal pathogens. These ExoY-like domains can be essential for virulence^13^. Elucidating the cellular activities and specificities of ExoY and ExoY-like toxins may therefore help to develop new therapeutic strategies against the toxicity and virulence of several pathogens.

Despite the progress in understanding downstream effects of ExoY activity, fundamental information on ExoY is lacking: as other bacterial soluble related cyclases such as CyaA and EF, ExoY is inactive in bacteria and is activated by an eukaryotic cofactor after its delivery to the target cells^5^. While the other class II adenylate cyclase toxins like CyaA and EF are strongly activated upon interaction with calmodulin^14, 15^, calmodulin is unable to stimulate ExoY enzymatic activity and the precise nature of the eukaryotic activator has remained elusive up to now. Here, we describe the identification of actin as the cofactor that activates *P. aeruginosa* ExoY and the ExoY-like module present in MARTX toxin of *V. nigripulchritudo* in host cells. Our findings suggest that, within the class II adenylyl cyclase family^16^, a new class of nucleotidyl cyclase toxins share actin as a common host activator.

## RESULTS

### An activator of ExoY is present in *Saccharomyces cerevisiae*

Arnoldo et al. have reported that overexpression of ExoY impairs yeast growth^17^, suggesting that ExoY should be catalytically active in this organism and therefore that a cofactor required for ExoY catalytic activity should be present in yeast. To test this hypothesis, we prepared extracts from *Saccharomyces cerevisiae* BY4741 cells and measured adenylate cyclase activity of recombinant ExoY carrying an N-terminal His-Flag tag (HF-ExoY) in the presence of increasing amounts of yeast cell extract *in vitro*. Extract from HeLa cells were used as control. We observed a dose dependent stimulation of ExoY activity by yeast cell extract, to levels that were similar to those measured when using the extract from HeLa cells (Fig. 1a). Thus, we decided to use *S. cerevisiae* as a convenient experimental system to identify the putative yeast activator that was likely to be evolutionarily conserved in human cells.

**Figure 1.**
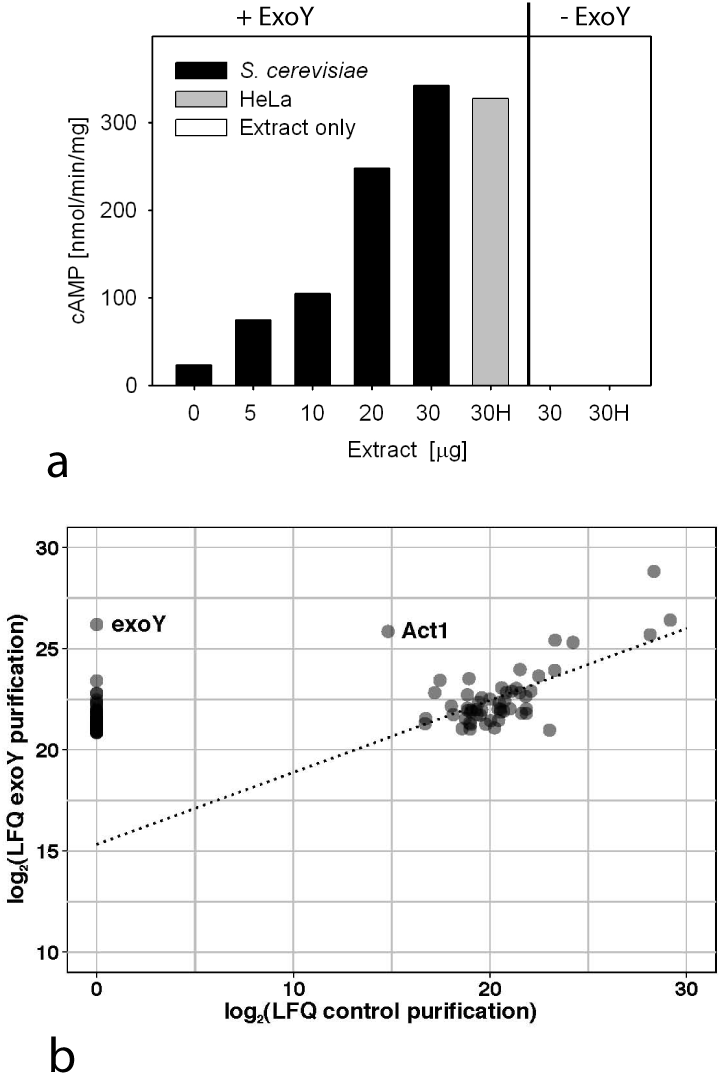
Presence of an activator of ExoY in *Saccharomyces cerevisiae*. **(a) Activation of HF-ExoY by extracts from HeLa cells or *S. cerevisiae***. 50 μl reactions containing 1 µg ExoY were started by the addition of the substrate ATP and stopped after 30 min incubation at 30 °C and the amount of synthesized cAMP was measured. **(b) Specific association of yeast Act1 to ExoY^K81M^**. Log_2_ transformed LFQ scores for the proteins identified in the fraction that copurified with ExoY^K81M^-TAP (y axis) were represented as a function of the scores obtained for the control purification (ExoY^K81M^-HA, x axis). Black circles are the result of two or more superimposed grey circles. For clarity, only the 100 proteins with highest LFQ scores in the TAP purification are shown; 45 of these factors, including ExoY, were not identified in the control purification and are represented on the y axis alone. The dashed line was computed by linear regression for the 55 proteins having LFQ values in both experiments and indicates the trend for common contaminants in the affinity purification.

### Identification of actin as the main protein interacting with ExoY-TAP in *S. cerevisiae*

To identify putative ExoY-binding proteins in yeast, we expressed ExoY containing a C-terminal epitope-tag (ExoY-TAP or ExoY-HA) in *S. cerevisiae* to isolate proteins co-purifying with the affinity purified bait protein. To avoid toxic effects due to cyclic nucleotide accumulation, we expressed a catalytically inactive variant of ExoY, ExoY^K81M^ (in which the Lys81 was changed to Met^5^). Material obtained in a one-step purification using IgGs covalently bound to magnetic beads was directly processed by tryptic digestion for protein identification through LC-MS/MS. The raw data were then analyzed by MaxQuant for protein identification and quantitative estimation of the specific enrichment of proteins in the experimental sample (ExoY^K81M^-TAP) as compared to the control (ExoY^K81M^-HA). While many abundant proteins were present in both samples to a comparable degree, as estimated from the label free quantitation score (LFQ,^18^), ExoY was identified only in the purification done with ExoY^K81M^-TAP extract, as expected. A second protein that was about 1000 times more abundant in the ExoY^K81M^-TAP purification than in the control was yeast actin (Uniprot P60010, YFL039C, Act1), which showed an LFQ score close to the score of the tagged ExoY (Fig. 1b). Other factors were identified specifically in the ExoY^K81M^-TAP purification, but with much lower LFQ scores (see Supplementary table 2). These results suggest a specific interaction of ExoY^K81M^ with yeast Act1. Since actin is one of the most highly conserved and abundant proteins in eukaryotic cells, it appeared to be a potential candidate for activating ExoY ubiquitously in eukaryotic cells.

### ExoY interacts with mammalian actin *in vitro*

To verify the interaction between ExoY and mammalian actin *in vitro*, we performed Ni-NTA agarose pulldowns. ExoY with a C-terminal Flag-His tag (ExoY-FH) and α-actin from rabbit skeletal muscle (Cytoskeleton, Inc., designated here MA-99) were added at equimolar concentrations to Ni-NTA agarose beads in batch. After 1 hr incubation the beads were washed in Durapore columns and the retained proteins were eluted with imidazole. While very little actin was bound unspecifically to the beads in the absence of ExoY, considerably more actin was present in the eluate from the sample containing ExoY-FH, suggesting specific binding between the two pure proteins (Fig. 2a).

**Figure 2.**
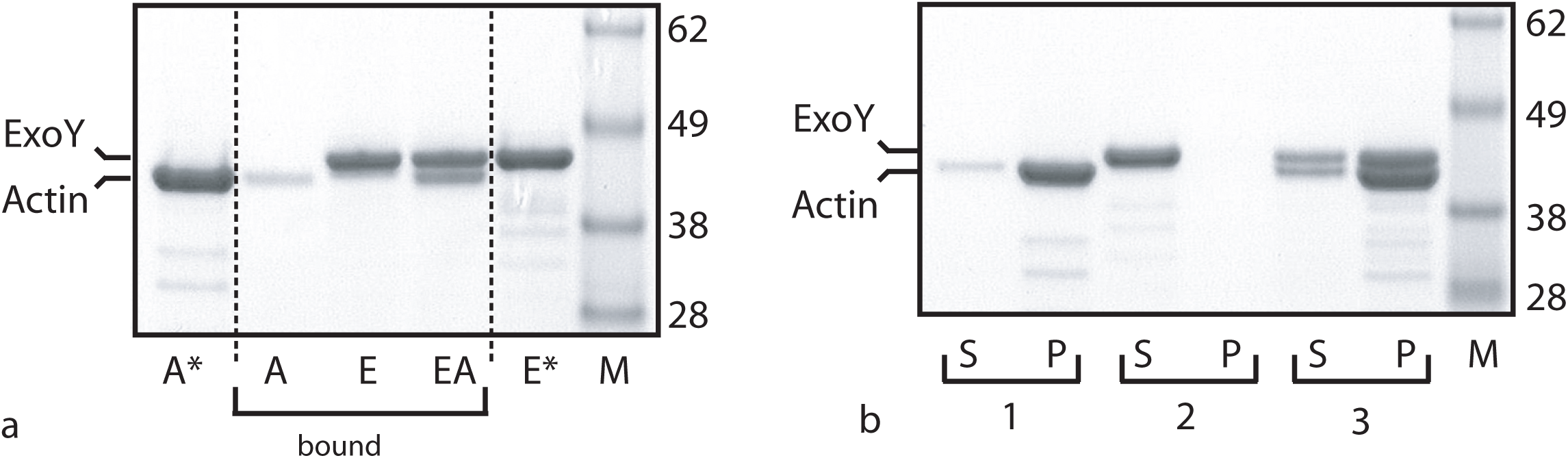
Interaction between ExoY and skeletal muscle actin from rabbit (MA-99). **(a)** 12.5 μg of actin (MA-99) (A), ExoY-FH (E), or each actin and ExoY-FH (EA) were allowed to bind to 5 μl Ni-NTA agarose in 450 μl binding buffer. Aliquots of immidazole-eluted fractions were separated on 4–12 % NuPAGE^®^ Bis-Tris gels (Invitrogen) in NuPAGE^®^ MES SDS running buffer and the gel was stained with Bio-Safe^TM^ Coomassie stain. Lanes A* and E* show the corresponding input for actin and ExoY alone, respectively. Corresponding amounts were loaded to allow direct comparability. Aliquots correspond to 15% of the total sample volume. The band of lower molecular weight was confirmed to be actin by western blots with anti-actin C4 antibodies. **(b) Co-sedimentation of ExoY with skeletal muscle F-actin**. Supernatant (S) and pellet (P) fractions were separated on 4–12 % NuPAGE^®^ Bis-Tris gels (Invitrogen) in NuPAGE^®^ MES SDS running buffer and the gel was stained as above. Reaction (1): actin only, (2): ExoY-FH only, (3) actin plus ExoY-FH.

Under our experimental conditions, which promote actin spontaneous polymerization (300 mM NaCl, 2.5 µM ATP, 660 nM actin), actin should exist in solution both in its monomeric (G-actin) and filamentous (F-actin) states. To investigate more specifically whether ExoY could bind to F-actin, we performed high-speed co-sedimentation assays: we polymerized G-actin-ATP and incubated F-actin at steady state subsequently with ExoY. Samples were then centrifuged at high speed to separate F-actin present in the pellet from G-actin present in the supernatant. Fig. 2b shows that ExoY, which alone partitioned in the supernatant fraction, was mostly found in the pellet fraction in the presence of F-actin, indicating ExoY capacity to interact with F-actin.

### Actin stimulates ExoY nucleotidyl cyclase activity *in vitro*

We next tested whether purified actin could activate ExoY *in vitro*. Highly pure non-muscle (cytoplasmic) actin isolated from human platelets (Cytoskeleton, Inc., designated here A-99) strongly stimulated the adenylate cyclase activity of ExoY (HF-ExoY, having 6xHis and Flag tags at the N-terminus), with a maximal activity reaching 120 µmol of cAMP.min^−1^.mg^−1^. Since in mammalian cells the transfection with a vector expressing ExoY led to accumulation of cGMP to levels exceeding that of cAMP^7, 8^, we also tested GTP as substrate and found that, in agreement with the preferential accumulation of cGMP over cAMP observed *in vivo*, the guanylate cyclase activity of HF-ExoY was approximately 8 times higher than the adenylate cyclase activity in the presence of actin *in vitro* (Fig. 3a). The background activity without actin was estimated to be about 1 nmol and 10 nmol.min^−1^.mg^−1^ for cGMP and cAMP, respectively. Thus, the ExoY nucleotidyl cyclase activity was stimulated more than 10,000 fold by submicromolar concentrations of F-actin. Different mammalian actin isoforms (A-99, a mixture of 85% β- and 15% γ-actin, or α-actin from rabbit skeletal muscle) led to a similar activity for cGMP synthesis (Supplementary Fig. 2). All together these data indicate that actin is a specific activator of ExoY in eukaryotic cells. Subsequent experiments were performed using highly pure skeletal muscle α-actin purified in one of our laboratory (designated MA-L).

**Figure 3.**
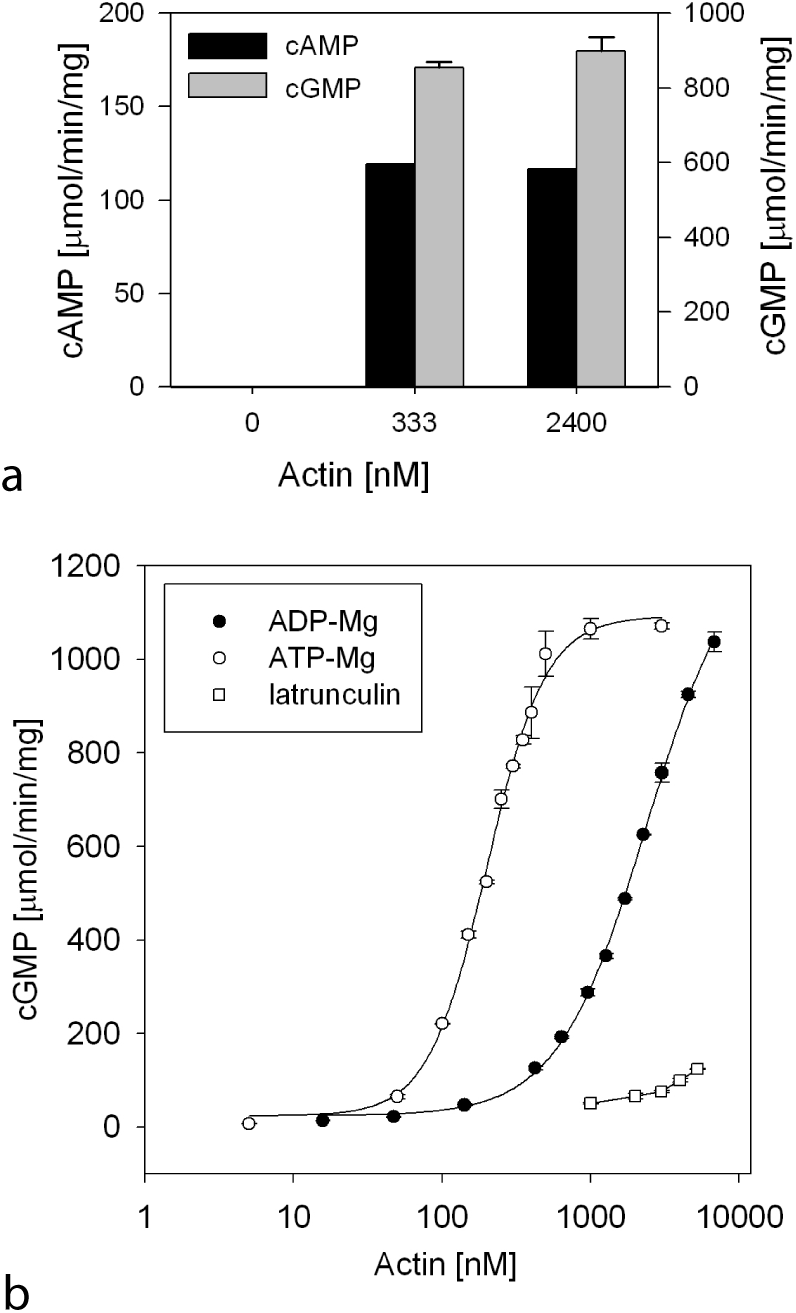
ExoY is efficiently activated by actin *in vitro*. **(a) Preferential synthesis of cGMP as compared to cAMP by ExoY activated by actin (A-99)**. Reactions containing HF-ExoY at 0.5 nM (1 ng) and actin at concentrations indicated were started by the addition of 2 mM ATP or GTP substrate and incubated for 10 min at 30 °C. **(b) Dependance of ExoY activation on the nucleotide and polymerization state of actin**. Muscle α-actin (MA-L) was converted to Mg-ATP-actin or Mg-ADP-actin and used at the concentration indicated to activate ExoY-FH. Mg-ATP-actin or Mg-ADP-actin polymerize above 100 and 1700 nM, respectively. Activities with the polymerization-inhibiting drug latrunculin A and Mg-ATP-actin (according to Figure 4) were plotted as comparison.

To examine a possible dependence of ExoY activation on the different states of actin (ATP- versus ADP-bound, monomeric versus polymeric forms), we measured ExoY cGMP synthesis activity at different actin concentrations below or above the critical polymerizing concentrations in different actin nucleotide states. Measurements were performed at various concentrations of actin that was initially loaded with either Mg-ATP or Mg-ADP. A similar maximal activity of 1000 - 1200 μmol of cGMP.min^−1^.mg^−1^ was obtained with both ATP-bound and ADP-bound actin (Fig. 3b). In contrast, the actin concentrations required for half maximal activation of ExoY (K_1/2_) were dependent upon the bound nucleotides. The actin concentration for half maximal ExoY activation with ATP-loaded actin was about 0.2 μM (Fig. 3b). This is just above the critical concentration of 0.1 μM above which Mg-ATP-actin spontaneously polymerizes in solution with salt^19^. Conversely, the actin concentrations required for half-maximal ExoY activation with ADP-loaded actin was about 2.4 μM (Fig. 3b), a shifted value that correlates with the alternative critical concentration of 1.7 obtained with Mg-ADP-actin. Altogether, these results suggest that the maximal activation of ExoY by actin was correlated with F-actin formation.

### Activation of ExoY by actin is antagonized by latrunculin A or G-actin binding proteins

We then examined whether proteins or molecules that are known to bind to G-actin and to inhibit its polymerization or spontaneous nucleation, could affect the activation of ExoY by actin. We examined the activation of ExoY by G-actin in the presence of the drug latrunculin A or G-actin binding proteins such as profilin or thymosin-β4 (Tβ4), which are among the main monomeric actin-binding proteins in vertebrate cells^19^. These three molecules are known to inhibit actin polymerization or spontaneous nucleation by binding to distinct G-actin interfaces. The small macrolide Latrunculin A from the Red Sea sponge *Negombata magnifica*^20, 21^ inhibits actin self-assembly by binding (*K_D_* ≈ 0.2 μM) to a cleft located on the pointed face of G-actin. The protein profilin binds (*K_D_* ≈ 0.1 μM) in contrast to the opposite face of monomers, called barbed face, and favors *in vivo* the unidirectional elongation of the most-dynamic barbed ends of filaments. *In vitro*, G-actin:profilin complexes inhibit actin spontaneous nucleation and thus polymerization in absence of actin nuclei or filament seeds. Tβ4 is a small intrinsically disordered β-thymosin domain of 4 kDa that acts as a major G-actin-sequestering polypeptide in cells^22, 23^. Here, we used a chimeric β-thymosin domain between Tβ4 and ciboulot from Drosophila called Chimera 2 (CH2), as it exhibits a higher affinity for G-actin than Tβ4 (*K_D_*~0.5 μM versus 2 μM) while retaining its sequestering activity^22, 23^. Like Tβ4, CH2 displays an extended binding interface on actin monomers by interacting with both their barbed and pointed faces^22, 23, 24^.

In ExoY activity measurements, actin monomers were saturated by the above molecules to inhibit or significantly slow down the spontaneous nucleation or polymerization of actin in solution. The inhibitory effect on actin assembly was verified in cosedimentation assays (Supplementary Fig. 3). In these conditions, all molecules tested inhibited ExoY activation by micromolar actin concentrations (Fig. 4), which normally induce maximal activation of the toxin. CH2 reduced at least 9 fold (up to 15) the ExoY activity measured in the presence of 1-3 μM of actin (Fig. 4). Latrunculin A reduced at least 7 fold (up to 13) and profilin decreased at least 4 fold (up to 7) the ExoY activity at similar ranges of actin concentrations (Fig. 4). Latrunculin A, profilin, and CH2 did not affect the low background activity of ExoY in the absence of actin. These data thus indicated that filamentous-actin is the preferred activator of ExoY.

**Figure 4.**
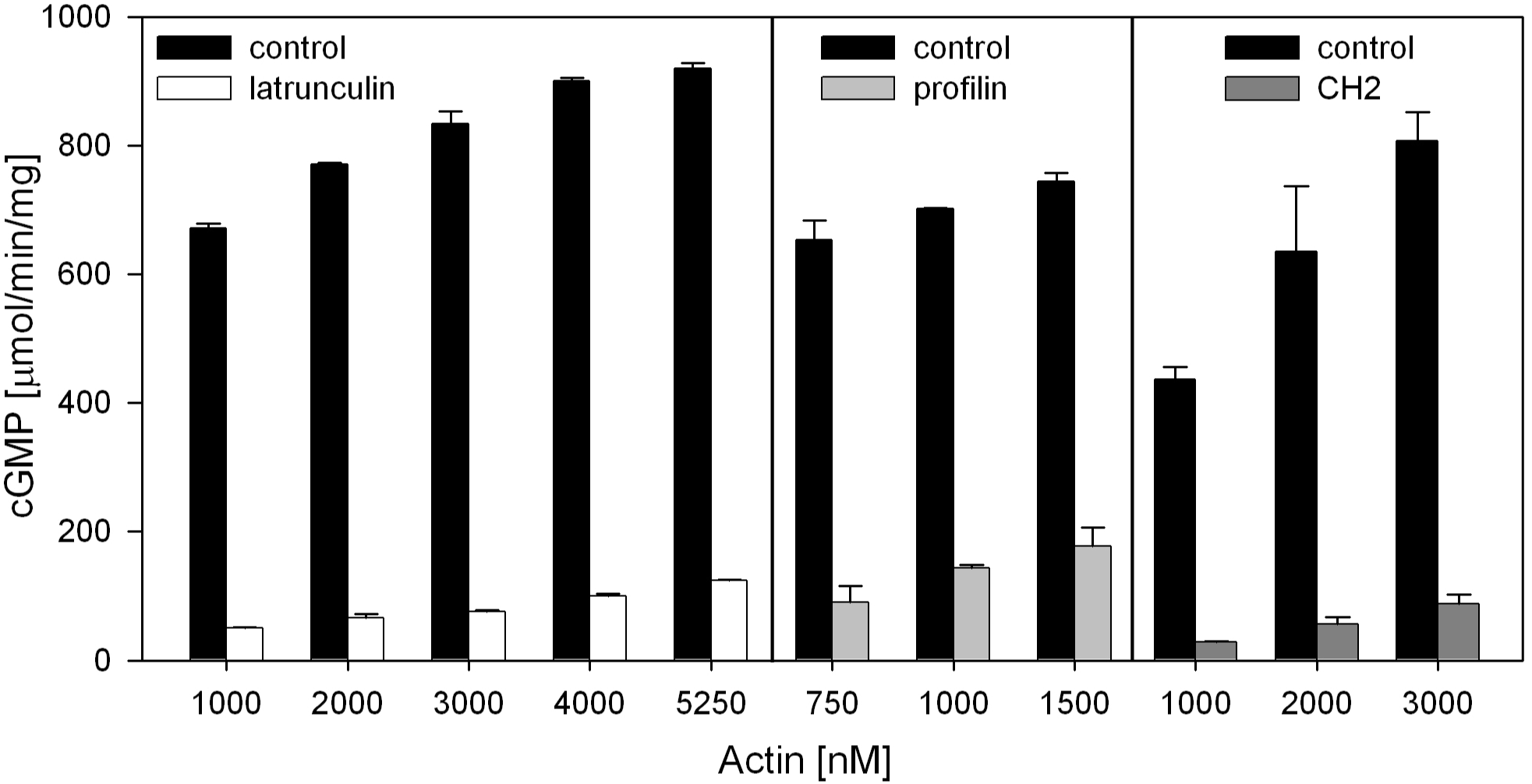
Effect of latrunculin A and G-actin binding proteins on the activation of ExoY. Reactions containing actin (+/- inhibitor) and ExoY-FH were preincubated for 10 min at 30 °C and started by the addition of 2 mM GTP and continued for 10 min. **Latrunculin A**: latrunculin A (at a final concentration of 10 μM) was added to actin and preincubated for 10 min at room temperature before conversion to Mg-ATP-actin and processing as for the control. Profilin: Profilin was added to Ca-ATP-G-actin at a 2:1 ratio. Control reactions lacking latrunculin or profilin contained G-actin (MA-L) that was converted to Mg-ATP-G-actin and allowed to polymerize to steady state conditions before dilution to the indicated concentrations. Chimera 2 (CH2) for a final concentration of 5 μM was added to muscle G-actin (MA-L) under conditions preventing salt-induced polymerization in G-buffer and preincubated for 10 min at 30 °C. None of the molecules tested affected the low basal ExoY activity in the absence of actin.

### ExoY is an F-actin binding protein that can modify the intrinsic or regulated dynamics of filaments by binding along filament sides

We next examined whether ExoY affects the intrinsic or regulated dynamics of actin self-assembly *in vitro* in assembly/disassembly assays with ExoY /ExoY^K81M^. The kinetics of polymerization or depolymerization were monitored by following the increase or decrease, respectively, in pyrene-actin fluorescence intensity (pyrenyl-labeled actin subunits exhibit higher fluorescence when incorporated in filaments than free in solution). In polymerization kinetics, ExoY slightly accelerated the rate of G-actin-ADP-Mg (Fig. 5a) and G-actin-ATP-Mg (Fig. 5b) self-assembly, confirming that ExoY can interact with actin without preventing its self-assembly. Yet, this stimulation of G-actin-ATP/ADP polymerization by ExoY was detectable only at high ExoY concentrations (in µM range). The dose-dependent acceleration was independent of the ExoY adenylate cyclase activity since it was observed with the inactive ExoY^K81M^ variant and also with the wild-type ExoY when only ADP was present. We further examined whether the ExoY-stimulation of actin polymerization was achieved by increasing elongation rates on barbed- or pointed-end, or by severing filaments, but found no effects of the toxin on these processes (Supplementary Fig. 4). We were unable to isolate stable G-actin/ExoY complexes in solution even at micromolar concentrations of both proteins (and in presence of latrunculin A to prevent actin polymerization). Besides, the ExoY-induced stimulation of actin polymerization was fully inhibited when actin was saturated by profilin (Fig. 5b). These results confirm that ExoY is unlikely to stimulate actin polymerization in host cells. Indeed, profilin-actin complexes form the major part of the polymerization competent G-actin pool within eukaryotic cells^19, 24^

**Figure 5.**
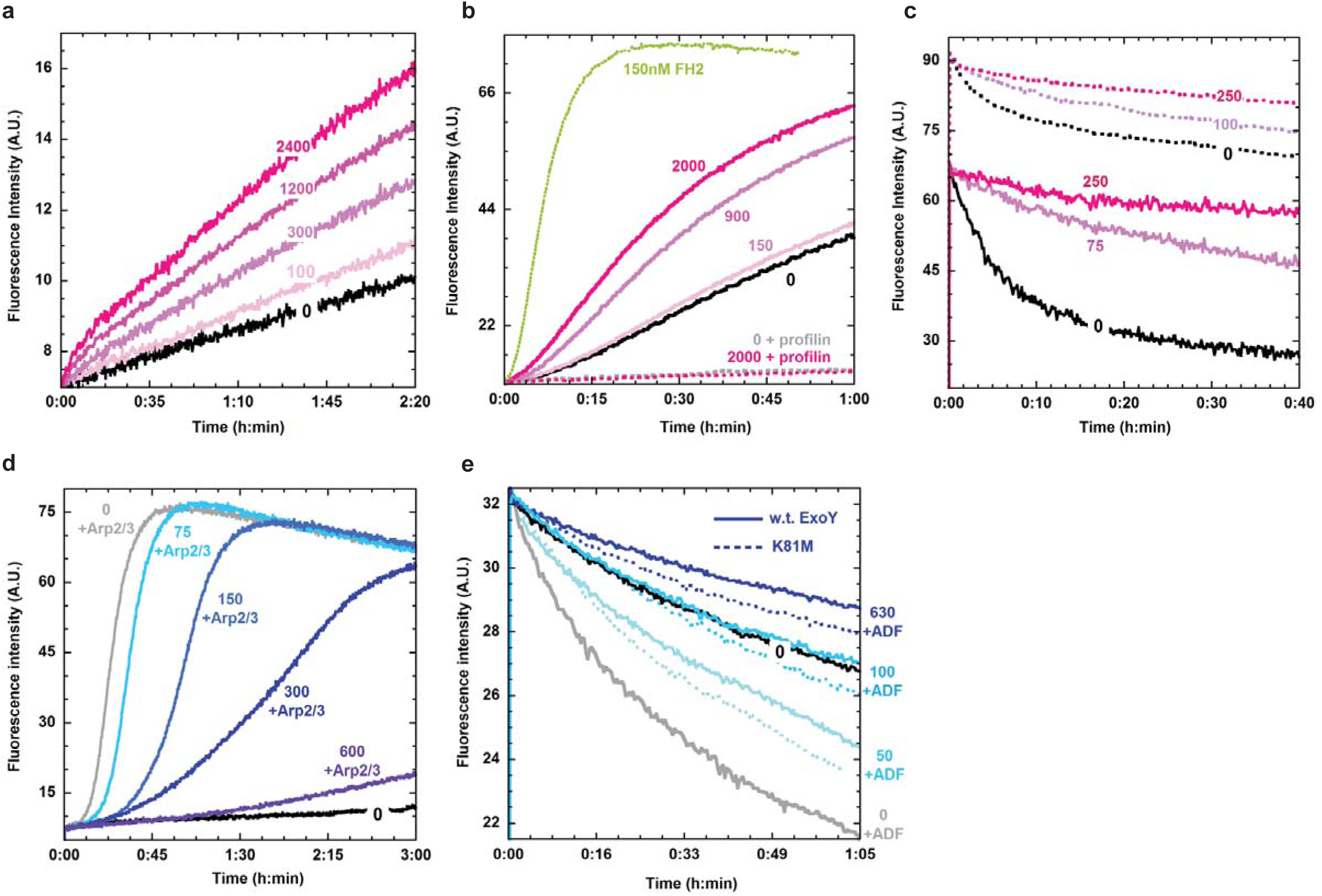
ExoY is an F-actin binding protein whose binding along the sides of filaments can interfere with the intrinsic or regulated dynamics of filaments. **(a) ExoY-ST mediated acceleration of G-actin-ADP-Mg self-assembly rate**. 8 μM G-actin-Mg-ADP (10% pyrenyl labeled) was polymerized in ExoY-ST absence or presence at the indicated concentrations (nM). **(b) ExoY^K81M^-ST mediated acceleration of G-actin-ATP-Mg selfassembly rate and its inhibition by profilin**. 4 μM G-actin-Mg-ATP (3% pyrenyl labeled) was polymerized in the absence (continuous lines) or presence (dashed lines) of profilin (at 10 μM) with 0 to 2000 nM ExoY^K81M^-ST. Efficient polymerization acceleration by a genuine G-actin nucleator, namely the dimeric FH2 domain of INF2 formin^51^ (in green, 50 nM) is shown for comparison. (c) Effects of ExoY**^K81M^**-ST binding to filaments on their spontaneous disassembly kinetics from free barbed (continuous lines) or pointed (dotted lines) ends. From free barbed-ends: F-actin (2 µM, 50% pyrenyl) at steady state was diluted to 50 nM in presence of 0 to 250 nM ExoY^K81M^-ST. Disassembly from pointed-ends: filaments with their barbed-ends capped by Gelsolin (10 µM actin, 33.3 nM Gelsolin) at steady state were diluted to 300 nM in presence of 0 to 250 nM ExoY^K81M^-ST. Different fluorescence intensity levels were used between the two depolymerization assays. **(d) Effect of ExoY^K81M^ -ST on the acceleration of filament formation induced by VCA-activated Arp2/3 complex**. 3 μM G-actin-Mg-ATP (3% pyrenyl labeled) was polymerized with 7.5 μM profilin, 0.2 μM NWASP-VCA, in absence (black) or presence of 35 nM Arp2/3 and 0 to 600 nM ExoY^K81M^-ST. **(e) Effect of ExoY/ExoY^K81M^-ST on the acceleration of F-actin- ADP disassembly promoted by ADF**. F-actin-ADP (9 μM) at steady state was diluted to 4 μM and preincubated for 2 min with 0 to 630 nM ExoY-ST (w.t., continuous lines) or ExoY^K81M^-ST (dotted lines) prior to depolymerization assays without ADF (0 nM ExoY/ExoY^K81M^, black) or with 4 μM ADF.

To delineate the interaction of ExoY with F-actin, we performed dilution-induced depolymerization assays monitoring filament disassembly from free barbed- and pointed ends. As shown in Fig.5c, ExoY^K81M^ inhibited the spontaneous disassembly of F-actin induced by dilution. This indicates that ExoY directly binds to filaments and thus stabilizes actin inter-subunit contacts. The inhibition of filament disassembly by ExoY^K81M^ was also observed when barbed ends were capped by gelsolin (Fig. 5c), thus excluding the possibility that ExoY inhibited disassembly by binding to barbed ends. These results and the absence of ExoY effects on the pointed-end elongation rate (Supplementary Fig. 4) indicate that ExoY binds along the sides of filaments where it likely interacts with several adjacent actin subunits thus stabilizing actin inter-subunit contacts and preventing spontaneous disassembly of filaments.

The binding of ExoY to filamentous actin was then quantified by co-sedimentation assays using F-actin steadily polymerized in the presence of ADP-BeF3^−^ (that mimics the filament transition state F-actin-ADP-Pi) in order to keep most actin firmly polymerized despite the efficient ATP-hydrolyzing activity of ExoY. To quantify more rigorously the ExoY bound to F-actin on SDS-PAGE gels in presence of high concentrations of F-actin (as ExoY and actin have close molecular weights), we used an ExoY protein fused to the Maltose-Binding-Protein (MBP), MBP-ExoY (note that MBP-ExoY has also a C-terminal Strep-Tag). As shown in Fig. S5, MBP-ExoY or MBP-ExoY^K81M^, which alone partitioned in the supernatant fraction, both pelleted with F-actin in a dose-dependent manner, and were almost completely found in the pellet fraction at concentrations above 25 μM F-actin-ADP-BeF_3_^-^, i.e. much below the F-actin concentrations in cells^25^. The estimated dissociation constants (Kd) of MBP-ExoY or MBP-ExoY^K81M^ for F-actin-ADP-BeF3^−^ (0.8±0.3μM and 2.5±0.4μM, respectively) (Supplementary Fig. 5), were in the same range as that of a number of eukaryotic cytoskeletal proteins that bind along filaments ^26, 27, 28^.

Finally, we analyzed whether ExoY binding to F-actin could interfere with the regulation of filament dynamics by eukaryotic cytoskeletal proteins that are known to bind along the sides of filaments. We considered two key regulatory proteins that are ubiquitous among eukaryotic cells: Arp2/3 complex and Actin-Depolymerizing Factor (ADF). The Arp2/3 complex, upon its activation by VCA domains of the WASP family proteins, can attach to the side of a pre-existing filament and catalyzes actin filament branching^26, 29^. ADF/cofilin proteins, present at micromolar concentrations in eukaryotic cells, bind cooperatively and preferentially to F-actin-ADP subunits along filaments (Kd~0.1μM), stimulating their disassembly and thus the turnover of actin filaments in cells^19, 26^.

We performed actin polymerization assays with G-actin saturated by profilin to simulate a more physiological context^19, 24, 30^. As shown in Fig. 5d, the acceleration of actin polymerization by Arp2/3 (25-35 nM), activated by N-WASP VCA domain, was significantly inhibited by high concentrations (≥100 nM) of ExoY^K81M^. This inhibition demonstrates that ExoY could antagonize the binding of the activated Arp2/3 complex along filaments and hence VCA-Arp2/3 regulation. In dilution-induced F-actin-ADP disassembly assays, ExoY^K81M^ (100 nM) were able to completely inhibit the acceleration of filament disassembly promoted by ADF (4 μM) (Fig. 5e). This inhibition suggests that ExoY (at submolar ratio with respect to ADF) could prevent the cooperative binding of ADF along F-actin-ADP and its disassembling activity.

### ExoY co-localizes with actin fibers in mouse NIH3T3 cells

To our knowledge, the localization of ExoY in eukaryotic cells was not previously reported and is of particular interest in light of its direct interaction and good affinity with naked F-actin *in vitro*. To avoid potential toxic effects of ExoY when expressed in eukaryotic cells, we examined the localization of the catalytically inactive variant ExoY^K81M^ fused to AcGFP, a monomeric green fluorescent protein. The ExoY^K81M^-AcGFP fusion protein was constitutively expressed from pUM518 under the control of the Pcmv ie promoter in transiently transfected NIH3T3 cells. Acti-stain^TM^ 555 fluorescent phalloidin (Cytoskeleton, Inc.) was used to visualize F-actin in cells. Unlike GFP alone, which showed a homogeneous cytoplasmic and partially nuclear localization, the signal for ExoY^K81M^-AcGFP partially colocalized *in vivo* with F-actin filaments (Fig. 6).

**Figure 6.**
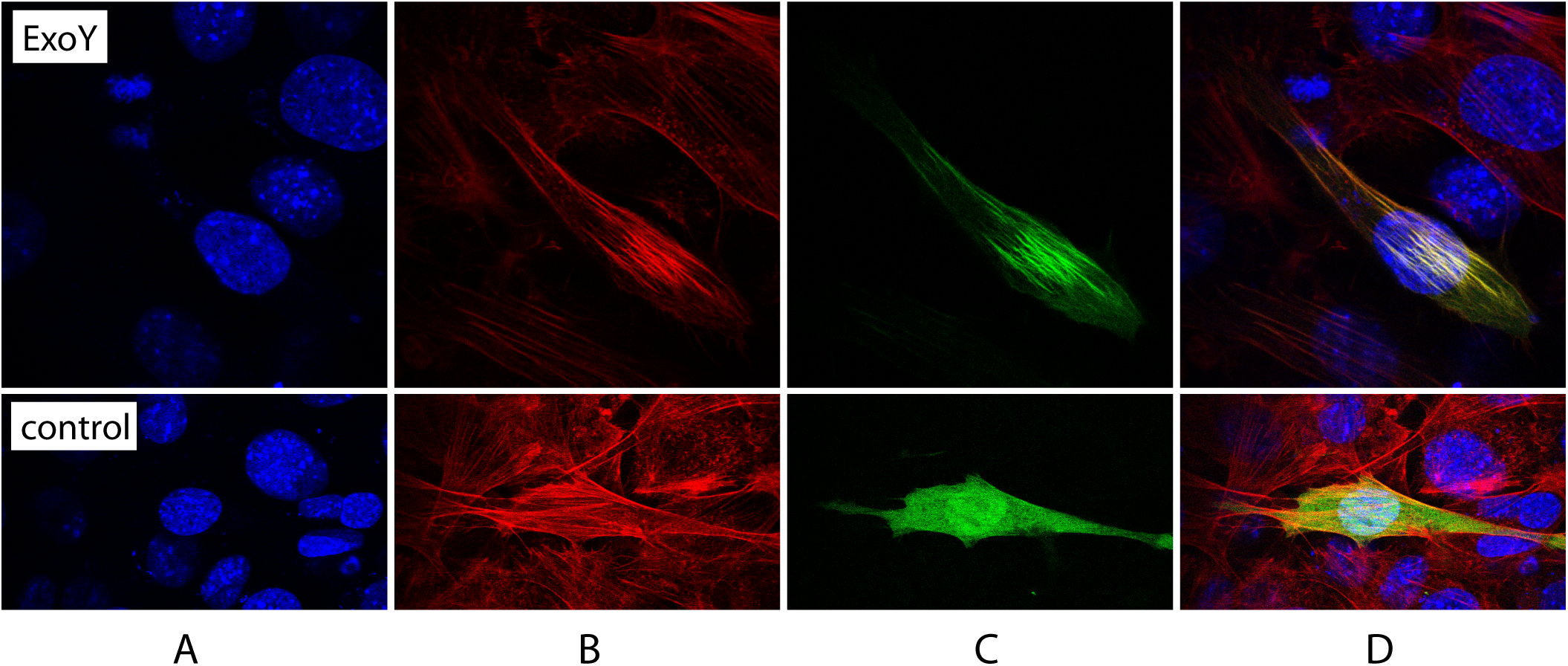
ExoY colocalizes with actin fibers in mouse NIH3T3 cells. Cells were transiently transfected with pUM518 expressing ExoY^K81M^-AcGFP (upper panel) or a control plasmid (pEGFP-C1) (lower panel) and stained with Acti-stain^TM^ 555 fluorescent phalloidin and DAPI. (A) nuclei stained with DAPI, (B) distribution of actin fibers stained with phalloidin, (C) distribution of ExoY^K81M^-AcGFP, and (D) merge of images (B) and (C).

### The ExoY-like virulence factor present in the *Vibrio nigripulchritudo* MARTX toxin is also activated by actin

Finally we examined whether actin could also activate other putative ExoY-like cyclases that can be found in various MARTX toxins that are produced by several Gram-negative pathogens including *Burkholderia* or several *Vibrio*^13^, *Providencia*, or *Proteus* species (Fig. 7a). For this, we selected the ExoY-like module from the MARTX toxin encoded by the virulence-associated plasmid pAs_Fn1_ from *Vibrio nigripulchritudo*^13, 31^, an emerging marine pathogen infecting farmed shrimps. The multidomain MARTX toxin is processed inside host cells by an inbuilt cysteine protein domain (CPD), into individual effector domains^32^. The N- and C-terminus of the *V. nigripulchritudo* ExoY-like effector domain were chosen based on sequence alignments of *P. aeruginosa* ExoY and ExoY-like containing sequences from several *Vibrio* MARTX toxins (Supplementary Fig. 6), the signature of CPD cleavage sites present in some *Vibrio* ExoY-like modules and HCA secondary prediction analysis. The MARTX-ExoY protein corresponding to residues Y3412 to L3872 of Uniprot reference F0V1C5 was termed here VnExoY-L. The protein carrying a C-terminal Flag-His tag (VnExoY-L-FH) was purified and tested for its adenylate cyclase activity in the presence and absence of actin. Fig. 7b shows that VnExoY-L displayed a potent adenylate cyclase activity in the presence of actin, which stimulates the enzymatic activity more than 10,000 fold. In contrast to *P. aeruginosa* ExoY, VnExoY-L did not exhibit any cGMP synthesizing activity. We conclude that actin may be a common activator of the various ExoY-like cyclase modules, even though these differ in their substrate selectivity.

**Figure 7.**
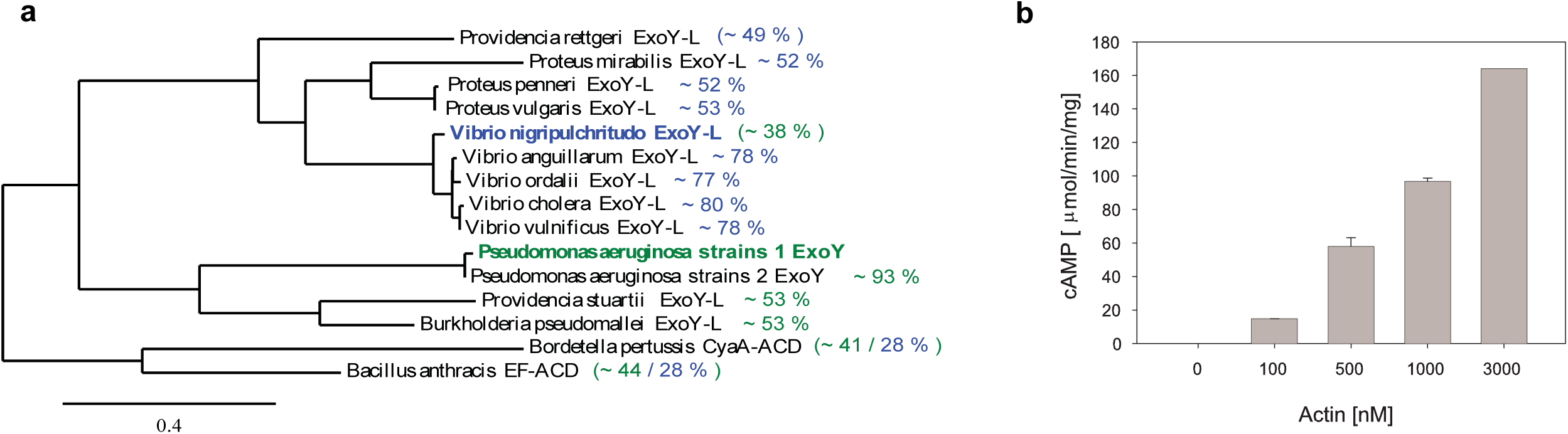
Phylogenetic tree of the bacterial ExoY-like nucleotidyl cyclase toxin subfamily (a) and activation of VnExoY-L catalyzed synthesis of cAMP by actin (b). **(a)** The amino-acid sequences of *Pseudomonas aeruginosa* ExoY and various ExoY-related effector domains/toxins found in several emerging Gram-negative bacterial pathogens were aligned as shown in Supplementary Fig. 6 and clustered on phylogram branches on the basis of the similarity of their amino acid sequences using the Phylogeny.fr platform^52^. Calmodulin-activated EF and Cya <u>A</u>denylate <u>C</u>yclase Domains (ACD) were used as an out group more distantly related to ExoY-like nucleotidyl cyclase toxins to root the phylogeny. NCBI accessions of bacterial protein sequences are given in Supplementary table 3. Pairwise sequence similarities (%) with *P. aeruginosa* ExoY or *V. nigripulchritudo* ExoY-like (VnExoY-L) are given in green and blue, respectively (to see supplementary table 3 for more details). Similarity values without parenthesis indicate the ExoY-like sequences that are the most significantly related to actin-activated ExoY or VnExoY-L nucleotidyl cyclases. **(b) Activation of VnExoY-L catalyzed synthesis of cAMP by actin (MA-L)**. Reactions containing VnExoY-L at 3.7 nM (10 ng) and actin at concentrations indicated were started by the addition of 2 mM ATP substrate and incubated for 30 min at 30 °C. The background activity without actin was estimated to be 1 nmol of cAMP.min^−1^.mg^−1^.

## DISCUSSION

Actin is the target of a variety of bacterial toxins. These toxins can affect the polymerization state of actin in different ways by introducing modifications, namely ADP-ribosylation at different position or crosslinking (for reviews see ^33, 34^). The provoked rearrangements have a profound effect on the cytoskeleton of the host cells and affect their response to bacterial invasion.

Here, we show that actin is a potent activator of a group of bacterial toxins that are homologous to the *P. aeruginosa* ExoY effector and that display nucleotidyl cyclase activities with different substrate selectivity.

We identified actin as a potential candidate for *P. aeruginosa* ExoY activation through its enriched presence among the proteins that co-purified with TAP-tagged ExoY expressed in *S. cerevisiae*. Actin is among the most abundant proteins in eukaryotic cells with a large number of known interaction partners. It is also frequently retrieved un-specifically in “pull-down” experiments and it is ranked among the top contaminants in the so-called “CRAPome”, Contaminant Repository for Affinity Purification^35^. In our case, however, we found that the interaction between ExoY and actin is highly specific and conserved between yeast and different mammalian actin isoforms. We further showed that ExoY is an F-actin binding protein (Fig. 5) and we demonstrated that F-actin is a potent activator of ExoY, able to stimulate its adenylate and guanylate cyclase activity more than 10,000 fold. In accordance with this view, we found that ExoY activation by actin was strongly antagonized by different G-actin binding proteins, such as profilin, or a Tβ4-derivative protein with similar activity as Tβ4 (CH2), or by latrunculin A that prevents actin polymerization (Fig. 4). Profilin and latrunculin A interact with non-overlapping, opposite binding sites on G-actin, which are mostly buried by actin:actin contacts in filaments. Profilin may partially overlap and compete with ExoY binding sites as it inhibits as well ExoY-mediated stimulation of actin self-assembly (Fig. 5b). In contrast, the antagonizing effect of the small molecule latrunculin A is likely due to its inhibition of actin polymerization rather than to a steric hindrance of the ExoY binding-sites. In the presence of saturating concentrations (> 2-5 μM) of F-actin, the very low basal enzymatic activity of ExoY was strongly stimulated to reach specific activities of about 120 µmol.min^−1^.mg^−1^ and 900 μmol.min^−1^.mg^−1^ for cAMP and cGMP synthesis, respectively. The higher guanylate cyclase activity as compared to the adenylate cyclase one is in agreement with the preferential accumulation of cGMP over cAMP observed *in vivo*^7, 8^. The corresponding kcat for cGMP synthesis is approaching 1000 s^−1^ and therefore within the same order of magnitude as the catalytic rates measured for cAMP synthesis for the related cyclase toxins CyaA or EF, when activated by calmodulin, their common eukaryotic activator^36, 37^

While ExoY represents to our knowledge the first example of a bacterial toxin that is activated by F-actin, G-actin has been shown before to activate a bacterial toxin secreted by the T3SS namely YopO/YpkA, a multidomain protein produced by pathogenic *Yersinia* species (Y. *enterocolitica* and *Y. pseudotuberculosis*, respectively), and involved in the disruption of the actin cytoskeleton^38^. YopO directly binds to an actin monomer and sterically blocks actin polymerization while, conversely, the bound actin induces autophosphorylation and activation of the YopO N-terminal serine/threonine kinase domain^39^. In this dimeric protein complex, the bound actin then serves as bait to recruit various host actin-regulating proteins that are then phosphorylated by YopO^40^. The mechanisms of activation of ExoY and YopO by F- and G-actin, respectively, are therefore likely different.

ExoY binds along naked filaments with a sub-micromolar affinity (Kd of about 0.8±0.3 μM, Supplementary Fig. 5) that should allow efficient competition *in vivo* with many eukaryotic cytoskeletal side-binding proteins that also bind along filaments with sub- to low micromolar affinities^26, 27, 28^. In agreement with this, we found that, *in vitro*, ExoY could antagonize the regulation of actin self-assembly dynamics by VCA-activated Arp2/3 complex (Fig. 5d) and/or the Actin-Depolymerizing Factor (ADF) (Fig. 5e): the enhanced turnover of actin filaments by ADF was inhibited by low and substoichiometric concentrations of ExoY. Considering the ExoY affinity for F-actin and the very high F-actin concentrations in eukaryotic cells (in non-muscle cells, G-and F-actin concentrations can reach 150 and 500 μM, respectively^25^), ExoY is expected to be fully bound to filaments within host cells, and this was indeed directly confirmed *in vivo* by the co-localization of ExoY-GFP with actin filaments in transfected NIH3T3 cells (Fig. 6). As many F-actin binding proteins, ExoY can also weakly interact with actin monomers as indicated by the fact that (i) ExoY is weakly activated (up to 5-10 % of maximal activity) in a dose dependent manner by actin bound to the polymerization-inhibiting drug latrunculin A (Figs. 3b and 4) and that (ii) ExoY weakly stimulated G-actin-ATP/-ADP polymerization in absence of profilin (Fig. 5a,b).

While most of the ExoY related effects in infected cells likely depend on its catalytic activity, the catalytically inactive ExoY^K81M^ mutant has been observed to induce temporary actin redistribution to the cell margins^10^ as well as minimal intercellular gap formation^11^ in endothelial cells. These effects may be linked to a residual nucleotide cyclase activity of ExoY^K81M^ and/or to its direct binding to actin polymers in host cells. We showed that *in vitro* (Fig. 5e), ExoY can antagonize the cooperative activity of ADF at sub-molar ratios of ExoY with respect to the regulatory cytoskeletal side binding protein. Although ExoY is likely present only at low concentrations in infected host cells, its binding along actin filaments could possibly contribute to the dysregulation of actin cytoskeleton dynamics *in vivo*. It will thus be interesting to examine in more detail the actin cytoskeleton dynamics of host cells upon infection with bacteria expressing catalytically inactive ExoY or ExoY-like proteins alone or together with other toxins affecting actin cytoskeleton regulation (ExoS, ExoT from *P. aeruginosa*, Actin Cross-linking (ACD) or Rho-GTPase Inactivation Domain (RID) from various MARTX toxins of the *Vibrio* genus^13^).

ExoY-like modules are frequently found among the effector domains of Multifunctional Autoprocessing RTX (MARTX) toxins in multiple bacterial species of the *Vibrio* genus^13^, which represent emerging human or animal pathogens. In addition, ExoY-like proteins can be found in various other Gram-negative pathogenic bacteria from the genus *Providencia, Burkholderia* or *Proteus* (Fig. 7a). Here we showed that VnExoY-L, a rather distantly related ExoY-like module from *V. nigripulchritudo* (38% sequence similarity with *P. aeruginosa* ExoY and 28% with *B. anthracis* EF or *B. pertussis* CyaA, Fig. 7a and Supplementary Table 3), was also strongly stimulated (over a 10,000 fold) by actin and efficiently synthesized cAMP but not cGMP. The lack of guanylate cyclase activity is in agreement with the results obtained with the *V. vulnificus* ExoY-like module^13^, a close homolog of VnExoY-L (>75% sequence similarity, Fig. 7a and Supplementary Table 3), and may thus reflect a more general difference regarding the nucleotide substrate specificities between the *P. aeruginosa* ExoY and other ExoY-like proteins found in MARTX toxins like those of the *Vibrio* genus (Fig. 7a and Supplementary Fig. 6).

Actin may thus represent a common eukaryotic activator for a sub-group (Fig. 7a) of the class II adenylyl cyclase toxin family (described in ^16^). This newly defined, actin-activated nucleotidyl cyclase (AA-NC) sub-family is also characterized by wider nucleotide substrate specificity than the original class II members, the adenylate cyclase toxins CyaA from *B. pertussis* and EF from *B. anthracis*. As calmodulin, the common cofactor of CyaA and EF, actin is an abundant and highly conserved protein specific to eukaryotic cells. It appears, therefore, to be a suitable molecular signal to indicate the arrival of the ExoY toxin in the eukaryotic environment of the host target cells, where it should display its cyclic nucleotide synthesizing activity.

Future studies should address the mechanism of activation of ExoY and ExoY-like proteins by actin through structural analysis. This could eventually open several interesting prospects in particular regarding the development of small molecules able to specifically inhibit the activation of these toxins by actin, as a therapeutic approach against bacterial infections, as well as the structural basis of the differential substrate selectivity of these AA- NCs.

Interestingly, Beckert et al.^7^ have reported notable differences in accumulation of various cNMPs in different cell lines exposed to *P. aeruginosa* ExoY, in particular, with respect to the mode of delivery of the toxin (transfection versus infection). It will be interesting to examine whether the relative efficacy in synthesizing different cNMPs depends solely on availability of substrates or whether the actin dynamics and turnover in cells may also play a role.

## MATERIAL AND METHODS

Strains, plasmids and growth conditions

Strains, plasmids and growth conditions are described in table S1.

### Purification of ExoY, actin from rabbit skeletal muscle, and actin-binding proteins

ExoY-FH and VnExoY-L-FH were purified by nickel affinity chromatography under denaturing conditions (in the presence of 8M urea) from the non-soluble protein fraction obtained from 1 liter cultures of *E. coli* BLR (pUM460) or (pUM522), respectively. Proteins were expressed from the λP*_L_*, promoter controlled by the temperature sensitive cI repressor (cI857), which was induced by shifting the temperature from 30°C to 42°C. Proteins were renatured by dialysis into 25 mM Tris pH8.0, 250 mM NaCl, 10 % glycerol, 1 mM DTT for ExoY-FH and 50 mM Tris pH 8.0, 200 mM NaCl, 10 % glycerol, 1 mM DTT for VnExoY-FH. HF-ExoY was purified from the soluble fraction obtained from 0.5 liter cultures of MG1655 (pUM447) that were grown at 30°C.

The fusion constructs of ExoY/ExoY^K81M^ with an N-terminal maltose-binding protein (MBP), designed as follows: (His-Tag)-(MBP)-(PreScission-site)-(ExoY/ExoY^K81M^)-(Strep-tagII) and referred in the text as MBP-ExoY/ExoY^K81M^-ST, or their truncated forms (ExoY/ExoY^K81M^-ST) were purified under non-denaturing conditions successively from HisTrap, StrepTrap, and Superdex 200 16*60 columns using standard protocols. The HisTag-MBP fusion was either cleaved or not using PreScission protease prior to the StrepTrap purification step. Proteins were stored in 25 mM Tris-HCl pH 8.8, 150 mM KCl, 3 mM NH4SO4, 0.5% Glycerol, 1 mM DTT.

Rabbit skeletal muscle alpha-actin was purified in our laboratory (refered as MAL) using several cycles of polymerization and depolymerization as previously described ^41^ and stored in G-buffer (5 mM Tris pH 7.8, 0.1 mM CaCl2, 0.2 mM ATP, 1 mM DTT). The purity of MA-L was estimated to be >95% according to analysis on denaturating SDS-PAGE gels. Functionality was controlled by several cycles of polymerization/depolymerization and by verifying that the measured actin concentrations of our samples fit the known critical concentration values of a fully functional and highly pure actin. ADP-actin was prepared by treatment of ATP-G-actin with hexokinase and glucose^42^.

Recombinant profilin I from mouse, chimera 2 of Thymosin-β4 and Drosophila ciboulot first β-Thymosin domain (CH2), full-length human gelsolin, VCA domain of human neural Wiskott-Aldrich syndrome protein (N-WASP), the Arp2/3 complex from bovine brain, or Spectrin-actin seeds from *human* erythrocytes were purified as described previously^22, 43, 44, 45^. The P529-P1013 FH2-WH2 construct of human INF2 formin (UniProt accession: Q27J81) was purified similarly as ExoY-ST.

### Affinity Purification

*S. cerevisiae* cells expressing ExoY_MUT_-TAP from pUM497 or ExoY_MUT_-HA from pUM498 as mock control were grown in 2 L YPGal at 30 °C. One step purification using Dynabeads^®^ magnetic beads conjugated to IgG were performed according to ^46^. One half of the methanol/chloroform precipitated protein was analyzed by PAGE followed by staining with Bio-Safe Coomassie G-250 (BIO-RAD) followed by silver staining (Pierce). The second half was directly digested by trypsin and analyzed by LC-MS/MS analysis at the proteomic facility of the Paris Descartes University (3P5) according to the details specified below. The raw data were analyzed by MaxQuant 1.3.0.5. software ^47^ for protein identification and quantitative estimation of the specific enrichment of proteins in the experimental sample as compared to the control.

### LC-MS/MS analysis

Proteomics analyses were realized at the 3P5 proteomics facility, Université Paris Descartes, Sorbonne Paris Cité, Institut Cochin, Paris as previously described^48^. Briefly: LC-MS protein analysis: peptides from Trypsin-digested extracts were concentrated, washed and analyzed using a reverse phase C18 column on an u3000 nanoHPLC hyphenated to a Linear Trap Quadrupole-Orbitrap mass spectrometer (all from Thermo). LTQ MS/MS CID spectra were acquired from up to 20 most abundant ions detected in the Orbitrap MS scan.

Protein identification: Proteome discoverer 1.3 (Thermo) with Mascot (matrixscience^49^) was used for protein identification. Separate analyses were compared using the MyPROMS software^50^.

### Interaction between ExoY and actin

12.5 μg of ExoY-FH, skeletal muscle actin from rabbit (AKL99, Cytoskeleton, Inc., designated in the text by MA-99), or both proteins in 450 μl of binding buffer [50 mM Na-phosphate pH 8.0, 300 mM NaCl, 20 mM imidazole, 0.01 % triton X-100, complete EDTA-free protease inhibitor cocktail (Roche)] were allowed to bind in batch to 5 μl Ni-NTA agarose (Quiagen) for 1 h at 4 °C rotating in Durapore filter units (Millipore). Unbound material was removed by centrifugation at 1,000 g for 1 min, after which the beads were washed three times with 450 μl of binding buffer and once with binding buffer supplemented 20 with 40 mM imidazole. Elution of bound proteins was performed by adding 50 μl of binding buffer supplemented with 0.5 M imidazole followed by incubation on ice for 10 min after which the eluate was collected by centrifugation at 3,000 g for 1 min. A second elution was carried out using 20 μl of elution buffer. A 12 μl aliquot of the combined eluates was analyzed by SDS-PAGE.

F-actin co-sedimentation assays (Fig. 2b) were performed using α-actin (MA-99) according to the instructions of Cytokeleton, Inc supplied with the “Actin binding Protein Biochem Kit^TM^ Muscle actin). 20 μl of a 48 μM actin (MA-99) solution in G’-buffer (5 mM Tris pH 8.0, 0.2 mM CaCl_2_, 0.5 mM DTT, 0.2 mM ATP, 5% glycerol) were thawed on ice, then added to 50 μl 50 mM Bis-tris propane (BTP) pH 9.5 and allowed to sit on ice for 40 min before polymerization was induced according to the protocol. 30 μl of polymerized F-actin stock solution were combined with 20 μl of a solution containing 12 μg ExoY-FH in 50 mM BTP pH 9.5, 270 mM NaCl, 2 mM DTT and 1x polymerization buffer from which non-soluble aggregates had been removed previously by centrifugation at 54,000 rpm in a TL55 rotor (Beckmann) at 18 °C for 1 h. This mixture as well as controls containing only actin or only ExoY and the corresponding buffers present in the experiment were incubated at room temperature for 30 min and centrifuged at 54,000 rpm for 90 min at 18 °C. Aliquotes of supernatant and resuspended pellet fraction corresponding to 15 % of the total samples were analyzed by SDS-PAGE.

To measure the equilibrium dissociation constant (Kd) of the ExoY:F-actin complex by co-sedimentation assays we used MBP-ExoY/ExoY^K81M^-ST and muscle α-actin (MA-L) (Supplementary Fig. 5). MBP-ExoY/ExoY^K81M^-ST should provide a reliable estimate of ExoY affinity for F-actin because these constructs perform similarily as ExoY/ExoY^K81M^-ST in depolymerization assays. MBP-ExoY/ExoY^K81M^-ST allowed separating and quantifying unambiguously by densitometry the fraction of the bound toxin at 88.9 kDa from actin at 42 kDa on SDS-PAGE gels, while ExoY-ST (M.W.of 43 kDa) was migrating too close to actin (M.W. of 42 kDa). No bundling activity was observed for ExoY in low-speed pelleting assays with F-actin. 1.5 μM of MBP-ExoY/ExoY^K81M^-ST was mixed for 1h with increasing amounts of F-actin-ADP-BeF3 (0 to 25 μM Mg-ADP-actin) at steady state in F1 buffer containing 5 mM ADP, 6 mM NaF, 0.6 mM BeCl2. The supernatant/unpolymerized (S) and pellet/polymerized (P) fractions were separated by an ultracentrifugation of 40 min at 200 000*g, resolved by 15% SDS-PAGE and detected by coomassie Blue staining. The ExoY-bound fraction was quantified by densitometry using the ImageJ software and plotted *versus* F-actin concentration. The following equation was used to fit the data, in which [E0] is the initial concentration of ExoY, [F0] the total concentration of F-actin in each measurement, and *K_d_* the equilibrium dissociation constant. The fraction *R* of ExoY bound to F-actin is as follows:

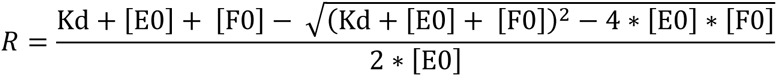

### Quantification of cAMP or cGMP synthesis *in vitro*

cAMP and cGMP synthesis were measured in 50 µl reactions containing 50 mM Tris pH 8.0, 7.5 mM MgCl_2_, 0.5 mg/ml BSA, 200 mM NaCl, 1 mM DTT, 2 mM ATP or GTP spiked with 0.1 μCi of [α-^33^P] ATP or [α^33^P] GTP, respectively, ExoY and indicated amounts of HeLa/S. *cerevisiae* cell extracts or purified actin (collectively termed activator). Reactions for Figs. 1A and 3A contained in addition 0.02% triton X-100, 0.1 mM CaCl_2_ and were lacking NaCl. Reactions were performed at 30 °C and were started by adding nucleotide substrate after a 5-10 minutes preincubation of ExoY plus activator. Under the conditions used, reactions were time linear for at least 20 minutes. Reactions were stopped by the addition of 450 μl stop solution (20 mM HEPES pH 7.5, 20 mM EDTA, 0.5 mM cAMP) and the mixtures were filtered on Al2O3 columns, which included 3 washes with 1 ml 20 mM HEPES pH 7.5 each to separate nucleotide substrates that were retained in the columns from cyclic nucleotides present in the filtrates. Filtrates were collected in 20 ml scintillation vials. 16 ml scintillation liquid (HiSafe3, Perkin Elmer) were added before measuring ^33^P in a TriCarb scintillation counter (Perkin Elmer). All reactions were performed in duplicates. Differences between cpm values of most duplicates were around or less than 10%. Standard deviations between duplicates are indicated by error bars.

Muscle actin 99% pure (designated MA-99) from rabbit skeletal muscle (Reference AKL99), or 99% pure non-muscle actin from human platelets (Reference APHL99, designated A-99) was obtained from Cytoskeleton, Inc. Alternatively, we used actin from rabbit skeletal muscle prepared in one of our laboratories (designated MA-L) according to the procedure described above. For activity assays, all actin solutions were diluted in G-buffer supplemented with BSA at 0.1 mg/ml.

Preliminary experiments to optimize reaction conditions showed that ExoY-FH was most active at pH values between 8 and 9 and in Tris as compared to HEPES or Na-phosphate and shows a broad optimal NaCl concentration (between 100 and 300 mM NaCl).

Extracts from HeLa cells for activation of ExoY were prepared as follows: Cells grown in Dulbecco's modified Eagle's medium (DMEM) + 10% fetal bovine serum were harvested after reaching 75% of confluence as follows: One wash with PBS was followed by incubation in 10 ml PBS containing 0.01M EDTA for 5 min at 37 °C before detaching the cells by gentle tapping of the flasks. Cells were collected by centrifugation and washed 3 times in PBS. The cell pellet was resuspended in 2 ml of lysis buffer [50 ml Tris pH 7.5, 300 mM NaCl, 0.5 % NP50, complete EDTA-free protease inhibitor cocktail (Roche)] per ml of cell pellet volume, after which the cells were snap frozen in liquid nitrogen and stored at -80 °C or processed immediately. Frozen cells were allowed to thaw on ice, rotated at 4 °C for 20 min and centrifuged for 1 h at 18,000 rpm in a SS34 rotor (Sorval). The supernatant was centrifuged at 100,000 rpm at 4 °C for 1 h in a TLA-110 rotor (Beckmann). The resulting supernatant was filtered through a 0.45 μM durapore PVDF filter unit (Millipore) and dialyzed against Tris/Triton/Glycerol buffer (TTG-buffer) containing 25 mM Tris pH 8.0, 0.1 % Triton X-100, 10 % glycerol, 1 mM DTT, 0.4 mM PMSF). Insoluble material was removed by centrifugation at 18,000 rpm in a SS34 rotor at 4 °C for 20 min. HeLa cell extract prepared according to this protocol contained approximately 10 mg/ml of protein and was stored in aliquots at -20 °C.

Extracts from *S. cerevisiae* were prepared from 100 ml cultures grown in YPD to an OD600 between 0.5 and 2, at which cells were harvested by centrifugation, washed once with water, and resuspended in 300 μl yeast lysis buffer [50 mM Tris pH 7.4, 50 mM KCl, 1 mM DTT, complete EDTA-free protease inhibitor cocktail (Roche)]. Cells were subsequently vortexed for a total of 5 min (5 times 1 min to prevent overheating) at 4 °C. Debris were removed by centrifugation at 16,000 rpm for 15 min at 4 °C. The resulting extract contained about 4 mg/ml protein and was stored at -20 °C after the addition of glycerol to a final concentration of 10 %.

Cycles of freezing and thawing did not seem to affect the activity of the cofactor necessary for ExoY activation in extracts from HeLa cells or *S. cerevisiae*.

Mg-ATP-actin was prepared from MA-L as follows: 90 μl of MA-L at 22.22 μM were added to 10 μl 10 x ME buffer (420 μM MgCl_2_, 4 mM EGTA) and incubated for 10 min at RT and put on ice.

A 34 μM solution of Mg-ADP-actin was prepared as follows: Ca-ATP-actin (MAL) was converted to Mg-ATP-actin by adding 100x concentrated ME-buffer and incubating for 10 min at RT to achieve a final concentration of 40 μM MgCl_2_ and 0.2 mM EDTA. 15 U/ml of Hexokinase (Roche) was added together with glucose to a concentration of 5 mM followed by 30 min incubation on ice. ApppppA was then added to 10 μM and the mixture was incubated for 5 min on ice. ADP and TCEP was added to 0.2 mM and 2 mM, respectively. Mg-ADP-actin was diluted in G-ADP-buffer (5 mM Tris pH 7.8, 0.2 mM ADP, 2 mM TCEP, 30 μM MgCl_2_). 15 μl of the diluted actin solutions (Mg-ATP-actin or Mg-ADP-actin) were combined with 30 μl of a mixture containing 1 ng ExoY in reaction buffer to achieve final reaction buffer conditions of 50 mM Tris pH 8.0, 200 mM NaCl, 7.5 mM MgCl_2_, 2 mM DTT and 0.5 mg/ml BSA.

Studies on the effect of profilin or latrunculin were performed with actin (MA-L) polymerized to steady state. For studies on the effect of profilin: (1) control reactions : 5 μl 5x concentrated ME-buffer (225 μM MgCl_2_, 2 mM EGTA) were added to 20 μl actin at 16.7 μM (diluted in G-buffer) yielding a concentration of 45 μM MgCl2 and 0.4 mM EGTA (25 ml Mg-ATP-actin (MA) and incubated for 10 min at room temperature. 25 µl fresh 2X polymerization buffer (F2: 300 mM KCl, 40 mM MgCl2, 10 mM ATP, and 10 mM DTT) was added and samples were allowed to sit at room temperature for 2 h to allow polymerization to proceed to steady state. Varying amounts of the so-prepared F-actin were combined with F1-buffer (F1: 150 mM KCl, 20 mM MgCl_2_, 5 mM ATP, and 5 mM DTT) to a total volume of 15 μl and added to 30 μl of a mixture containing 1 ng ExoY in buffer to achieve final reaction buffer conditions of 50 mM Tris pH 8.0, 200 mM NaCl, 7.5 mM MgCl_2_, 2 mM DTT and 0.5 mg/ml BSA. After 30 min preincubation at 30 °C, reactions were started by the addition of GTP (2 mM, spiked with [α-^33^P] GTP) and allowed to proceed for 10 min. (2) Reactions containing profilin: profilin was dialyzed against G-buffer to remove the KCl present in the storage buffer before adding 5 μl at 142.7 μM directly to undiluted actin (6 μl MA at 55.88 in G-buffer), incubated at room temperature for 10 min and diluted by adding 14 μl G-buffer. Actin was not converted into Mg-ATP-actin. 25 μl fresh F2 buffer was added and 11.2, 7.5, or 5.6 μl of this mixture were combined with G-buffer to a total volume of 15 μl and used immediately in activity assays. At final actin concentrations of 1.5, 1 and 0.75 μM, profilin was present at 3, 2, and 1.5 μM, respectively.

Latrunculin A was purchased from tebu-bio (produced by Focus Biomolecules). Studies with latrunculin were done similarly to those on profilin except that higher concentrations of actin (between 5.25 and 1.0 μM final MA-L) were used, ME buffer (10 fold concentrated) was added to control reactions before polymerization but was added as well to reactions containing latrunculin after combining latrunculin and actin and 10 min incubation at room temperature. Samples containing latrunculin were incubated 2 hours at room temperature as the corresponding controls. The mixture of lartunculin and actin was diluted to different concentrations in G-buffer containing latrunculin to ensure a finale concentration of 10 μM in all reactions. Control reactions lacking latrunculin contained DMSO at concentrations equivalent to that introduced with latrunculin.

Sedimentation-assays to verify effectiveness of profilin or latrunculin in preventing polymerization were performed at 2 or 1 μM actin (MA-L) in the exact same way as for measuring activity except that GTP was not spiked with [α-^33^P] GTP.

### Pyren-actin polymerization and depolymerization assays

Actin polymerization or depolymerization were monitored at 25°C by the increase or decrease in fluorescence, respectively, of 3-10% (polymerization) or 50% (depolymerization) pyrenyl-labeled actin (λexc = 340 nm, λem = 407 nm). Actin-Ca-ATP in G-buffer was converted just prior to the experiments into G-actin-Mg-ATP by adding 1/100 (vol./vol.) of (2 mM MgCl2, 20 mM EGTA). Polymerization or depolymerization were induced in a final F1-buffer containing (0.1 M KCl, 8 mM MgCl_2_, 50 mM Tris-HCl pH 7.8, 9 mM TCEP, 1 mM DTT, 0.3 mM NH4SO4), 10 to 15 mM ATP or ADP, and 3 to 5 mM GTP, unless indicated otherwise in figure legends. Fluorescence measurements were carried out in a Safas Xenius model FLX (Safas, Monaco) spectrophotometer, using a multiple sampler device. Dilution-induced depolymerization assays were performed by quickly diluting 4 to 68 μL of 9 to 13 μM 50% pyrenyl-labeled F-actin at steady state into a final volume of 160 μl containing F1 buffer, ATP and the proteins of interest.

### Localization of ExoY-AcGFP and actin in mouse NIH3T3 cells

NIH3T3 cells were grown in Dulbecco's modified Eagle's medium with high glucose (4.5 g/l) and 10% fetal bovine serum (FCS, PAA). Cells were plated on coverslips and transiently transfected by standard calcium phosphate precipitation the following day. Two days after the transfection, the cells were washed twice with cold PBS and fixed by 4% formaldehyde for 10 min at room temperature. Cells were then washed with PBS at room temperature, permeabilized with 0.5% Triton X100 in PBS and washed again with PBS. 200 μl of 100 nM Acti-stain^TM^ 555 fluorescent phalloidin (PHDH1, Cytoskeleton, Inc.) were added and the coverslip was incubated at room temperature for 30 min. Cover slips were mounted on glass slides using Fluoroshield mounting medium with DAPI (ab104139, Abcam). Cells were visualized with a confocal scanning microscope (LEICA SPE).

## ACKNOWLEDGEMENT

We thank Alain Jacquier and Micheline Fromont-Racine for support and discussions. We thank Alexander Ishchenko for providing us with a first yeast extract, Scot Ouellette for growing HeLa cells, Romé Voulhoux for chromosomal DNA of *P. aeruginosa*, Didier Mazel and Evelyne Krin for strain *V. nigripulchritudo*, the 3P5 proteomic facility, Emilie Cochet at the Institut Gustave Roussy for mass spectrometry analysis and Alexandre Chenal for help in data analysis. This project was funded by the Institut Pasteur under #PTR425 to UM and CS.

## REFERENCES

1. Engel J, Balachandran P. Role of Pseudomonas aeruginosa type III effectors in disease. Curr Opin Microbiol 12, 61–66 (2009).

2. Hauser AR. The type III secretion system of Pseudomonas aeruginosa: infection by injection. Nat Rev Microbiol 7, 654–665 (2009).

3. Burstein D, et al. Novel type III effectors in Pseudomonas aeruginosa. MBio 6, e00161 (2015).

4. Feltman H, Schulert G, Khan S, Jain M, Peterson L, Hauser AR. Prevalence of type III secretion genes in clinical and environmental isolates of Pseudomonas aeruginosa. Microbiology 147, 2659–2669 (2001).

5. Yahr TL, Vallis AJ, Hancock MK, Barbieri JT, Frank DW. ExoY, an adenylate cyclase secreted by the Pseudomonas aeruginosa type III system. Proc Natl Acad Sci USA 95, 13899–13904 (1998).

6. Gottle M, et al. Cytidylyl and uridylyl cyclase activity of bacillus anthracis edema factor and Bordetella pertussis CyaA. Biochemistry 49, 5494–5503 (2010).

7. Beckert U, et al. ExoY from Pseudomonas aeruginosa is a nucleotidyl cyclase with preference for cGMP and cUMP formation. Biochem Biophys Res Commun 450, 870–874 (2014).

8. Ochoa CD, Alexeyev M, Pastukh V, Balczon R, Stevens T. Pseudomonas aeruginosa exotoxin Y is a promiscuous cyclase that increases endothelial tau phosphorylation and permeability. J Biol Chem 287, 25407–25418 (2012).

9. Balczon R et al. Pseudomonas aeruginosa exotoxin Y-mediated tau hyperphosphorylation impairs microtubule assembly in pulmonary microvascular endothelial cells. PLoS One 8, e74343 (2013).

10. Cowell BA, Evans DJ, Fleiszig SM. Actin cytoskeleton disruption by ExoY and its effects on Pseudomonas aeruginosa invasion. FEMS Microbiol Lett 250, 71–76 (2005).

11. Sayner SL, Frank DW, King J, Chen H, VandeWaa J, Stevens T. Paradoxical cAMP-induced lung endothelial hyperpermeability revealed by Pseudomonas aeruginosa ExoY. Circ Res 95, 196–203 (2004).

12. Stevens TC, et al. The Pseudomonas aeruginosa exoenzyme Y impairs endothelial cell proliferation and vascular repair following lung injury. Am J Physiol Lung Cell Mol Physiol, (2014).

13. Ziolo KJ, Jeong HG, Kwak JS, Yang S, Lavker RM, Satchell KJ. Vibrio vulnificus biotype 3 multifunctional autoprocessing RTX toxin is an adenylate cyclase toxin essential for virulence in mice. Infect Immun 82, 2148–2157 (2014).

14. Leppla SH. Anthrax toxin edema factor: a bacterial adenylate cyclase that increases cyclic AMP concentrations of eukaryotic cells. Proc Natl Acad Sci U S A 79, 3162–3166 (1982).

15. Berkowitz SA, Goldhammer AR, Hewlett EL, Wolff J. Activation of prokaryotic adenylate cyclase by calmodulin. Ann N YAcad Sci 356, 360 (1980).

16. Barzu O, Danchin A. Adenylyl cyclases: a heterogeneous class of ATP-utilizing enzymes. Prog Nucleic Acid Res Mol Biol 49, 241–283 (1994).

17. Arnoldo A, et al. Identification of small molecule inhibitors of Pseudomonas aeruginosa exoenzyme S using a yeast phenotypic screen. PLoS Genet 4, e1000005 (2008).

18. Luber CA, et al. Quantitative proteomics reveals subset-specific viral recognition in dendritic cells. Immunity 32, 279–289 (2010).

19. Pollard TD, Blanchoin L, Mullins RD. Molecular mechanisms controlling actin filament dynamics in nonmuscle cells. Annu Rev Biophys Biomol Struct 29, 545–576 (2000).

20. Coue M, Brenner SL, Spector I, Korn ED. Inhibition of actin polymerization by latrunculin A. FEBS Lett 213, 316–318 (1987).

21. Yarmola EG, Somasundaram T, Boring TA, Spector I, Bubb MR. Actin-latrunculin A structure and function. Differential modulation of actin-binding protein function by latrunculin A. J Biol Chem 275, 28120–28127 (2000).

22. Didry D, et al. How a single residue in individual beta-thymosin/WH2 domains controls their functions in actin assembly. Embo J 31, 1000–1013 (2012).

23. Husson C, et al. Multifunctionality of the beta-thymosin/WH2 module: G-actin sequestration, actin filament growth, nucleation, and severing. Ann N Y Acad Sci 1194, 44–52 (2010).

24. Xue B, Leyrat C, Grimes JM, Robinson RC. Structural basis of thymosin beta4/profilin exchange leading to actin filament polymerization. Proc Natl Acad Sci U S A 111, E4596–4605 (2014).

25. Koestler SA, Rottner K, Lai F, Block J, Vinzenz M, Small JV. F- and G-actin concentrations in lamellipodia of moving cells. PLoS One 4, e4810 (2009).

26. dos Remedios CG, et al. Actin binding proteins: regulation of cytoskeletal microfilaments. Physiol Rev 83, 433–473 (2003).

27. Olshina MA, et al. Plasmodium falciparum coronin organizes arrays of parallel actin filaments potentially guiding directional motility in invasive malaria parasites. Malar J 14, 280 (2015).

28. Bugyi B, Didry D, Carlier MF. How tropomyosin regulates lamellipodial actin based motility: a combined biochemical and reconstituted motility approach. EMBO J 29, 14–26 (2010).

29. Ti SC, Jurgenson CT, Nolen BJ, Pollard TD. Structural and biochemical characterization of two binding sites for nucleation-promoting factor WASp-VCA on Arp2/3 complex. Proc Natl Acad Sci U S A 108, E463–471 (2011).

30. Rotty JD, et al. Profilin-1 serves as a gatekeeper for actin assembly by Arp2/3- dependent and -independent pathways. Dev Cell 32, 54–67 (2015).

31. Goudenege D, et al. Comparative genomics of pathogenic lineages of Vibrio nigripulchritudo identifies virulence-associated traits. Isme J 7, 1985–1996 (2013).

32. Shen A, et al. Mechanistic and structural insights into the proteolytic activation of Vibrio cholerae MARTX toxin. Nat Chem Biol 5, 469–478 (2009).

33. Aktories K, Lang AE, Schwan C, Mannherz HG. Actin as target for modification by bacterial protein toxins. Febs J 278, 4526–4543 (2011).

34. Haglund CM, Welch MD. Pathogens and polymers: microbe-host interactions illuminate the cytoskeleton. J Cell Biol 195, 7–17 (2011).

35. Mellacheruvu D, et al. The CRAPome: a contaminant repository for affinity purification-mass spectrometry data. Nat Methods 10, 730–736 (2013).

36. Drum CL, et al. Structural basis for the activation of anthrax adenylyl cyclase exotoxin by calmodulin. Nature 415, 396–402 (2002).

37. Karst JC, Sotomayor Perez AC, Guijarro JI, Raynal B, Chenal A, Ladant D. Calmodulin-induced conformational and hydrodynamic changes in the catalytic domain of Bordetella pertussis adenylate cyclase toxin. Biochemistry 49, 318–328 (2010).

38. Juris SJ, Rudolph AE, Huddler D, Orth K, Dixon JE. A distinctive role for the Yersinia protein kinase: actin binding, kinase activation, and cytoskeleton disruption. Proc Natl Acad Sci U S A 97, 9431–9436 (2000).

39. Trasak C, et al Yersinia protein kinase YopO is activated by a novel G-actin binding process. J Biol Chem 282, 2268–2277 (2007).

40. Lee WL, Grimes JM, Robinson RC. Yersinia effector YopO uses actin as bait to phosphorylate proteins that regulate actin polymerization. Nat Struct Mol Biol 22, 248–255 (2015).

41. Spudich JA, Watt S. The regulation of rabbit skeletal muscle contraction. I. Biochemical studies of the interaction of the tropomyosin-troponin complex with actin and the proteolytic fragments of myosin. J Biol Chem 246, 4866–4871 (1971).

42. Mizuno H, Higashida C, Yuan Y, Ishizaki T, Narumiya S, Watanabe N. Rotational movement of the formin mDia1 along the double helical strand of an actin filament. Science 331, 80–83 (2014).

43. Casella JF, Maack DJ, Lin S. Purification and initial characterization of a protein from skeletal muscle that caps the barbed ends of actin filaments. J Biol Chem 261, 10915–10921 (1986).

44. Gaucher JF, Mauge C, Didry D, Guichard B, Renault L, Carlier MF. Interactions of isolated C-terminal fragments of neural Wiskott-Aldrich syndrome protein (N-WASP) with actin and Arp2/3 complex. J Biol Chem 287, 34646–34659 (2012).

45. Le Clainche C, Carlier MF. Actin-based motility assay. Curr Protoc Cell Biol Chapter 12, Unit 12 17 (2004).

46. Defenouillere Q, et al. Cdc48-associated complex bound to 60S particles is required for the clearance of aberrant translation products. Proc Natl Acad Sci U S A 110, 5046–5051 (2013).

47. Cox J, Mann M. MaxQuant enables high peptide identification rates, individualized p.p.b.-range mass accuracies and proteome-wide protein quantification. Nat Biotechnol 26, 1367–1372 (2008).

48. Vallabhaneni KC, et al. Extracellular vesicles from bone marrow mesenchymal stem/stromal cells transport tumor regulatory microRNA, proteins, and metabolites. Oncotarget 6, 4953–4967 (2015).

49. Perkins DN, Pappin DJ, Creasy DM, Cottrell JS. Probability-based protein identification by searching sequence databases using mass spectrometry data. Electrophoresis 20, 3551–3567 (1999).

50. Poullet P, Carpentier S, Barillot E. myProMS, a web server for management and validation of mass spectrometry-based proteomic data. Proteomics 7, 2553–2556 (2007).

51. Chhabra ES, Higgs HN. INF2 Is a WASP homology 2 motif-containing formin that severs actin filaments and accelerates both polymerization and depolymerization. J Biol Chem 281, 26754–26767 (2006).

52. Dereeper A, et al. Phylogeny.fr: robust phylogenetic analysis for the non-specialist. Nucleic Acids Res 36, W465–469 (2008).

